# Deletion of CD73 increases exercise power in mice

**DOI:** 10.1101/2021.02.17.431631

**Authors:** Aderbal S Aguiar, Ana Elisa Speck, Paula M. Canas, Rodrigo A. Cunha

## Abstract

Ecto-5’-nucleotidase or CD73 is the main source of extracellular adenosine involved in the activation of adenosine A_2A_ receptors, responsible for the ergogenic effects of caffeine. We now investigated the role of CD73 in exercise by comparing female wild-type (WT) and CD73 knockout (KO) mice in a treadmill graded test to evaluate running power, oxygen uptake 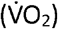, and respiratory exchange ratio (RER) – the gold standards characterizing physical performance. Spontaneous locomotion in the open field and submaximal running power and 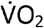 in the treadmill were similar between CD73-KO and WT mice; 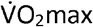 also demonstrated equivalent aerobic power, but CD73-KO mice displayed a 43.7±4.2% larger critical power (large effect size, P<0.05) and 3.8±0.4% increase of maximum RER (small effect size, P<0.05). Thus, KO of CD73 was ergogenic, i.e., it increased physical performance.

## INTRODUCTION

Adenosine is an inter-cellular modulator that signals altered cellular activity and metabolic stress within a tissue by activating adenosine receptors [1]. Acute exercise increases adenosine levels in human blood [2,3] and the rat brain [4]. Accordingly, adenosine contributes to exercise-induced vasodilation [5] and also causes drowsiness and tiredness at rest, being a candidate molecule to signal exercise fatigue [6–8]. The non-selective adenosine receptor antagonist, caffeine, is well established to cause an ergogenic effect [8,9], which is mimicked by selective antagonists of adenosine A_2A_ receptors (A_2A_R), abrogated in forebrain A_2A_R knockout mice [8], and eliminated upon administration of the A_2_R agonist NECA [6,10]. These observations strongly imply central A_2A_R as critical regulators of the impact of adenosine on exercise performance, whereas there is no consistent evidence for the involvement of A_1_, A_2B_, or A_3_ receptors in the control of exercise performance.

The activation of A_2A_R is achieved by a particular pool of extracellular adenosine formed by ecto-5’-nucleotidase, or CD73, responsible for the final formation of ATP-derived extracellular adenosine. In fact, the pharmacological or genetic inhibition of CD73 has effects identical to the inhibition of A_2A_R in the control of brain functioning [11–13], of the immune system [14] or adaptive vascular control in the periphery [15]. However, the role of CD73 in exercise remains to be defined, which prompted the present study to characterize the impact of deleting CD73 on the exercise performance of mice.

## METHODS

### Animals

Since we previously characterized the discrete impact of the estrous cycle on exercise performance of mice [16], we used 17 female mice (20.8±0.2 g, 8-10 weeks old) from our inbred colony with CD73-KO mice and wild-type (WT) littermates, cross-bred as previously described [13,17]. Mice were housed in collective home cages (n=3-5) under a controlled environment (12 h light-dark cycle, lights on at 7 AM, and room temperature of 22±1°C) with *ad libitum* access to food and water. WT and CD73-KO mice were housed together, with no genotype separation, following European Union guidelines (2010/63), and approval by the Ethical Committee of the Center for Neuroscience and Cell Biology (University of Coimbra, ORBEA 138-2016/1507201).

Mice were habituated to handling and the treadmill (9 m/min) in the three days before starting the experiments, which were performed between 9 AM and 5 PM, within the light phase of the dark/light cycle, in a sound-attenuated and temperature (20.3±0.6 °C) and humidity (62.8±0.4%) controlled room, under low-intensity light (≈ 10 lux). The open field apparatus and the treadmill were cleaned with 10% ethanol between individual experiments. For each test, the experimental unit was an individual animal.

### Open field

Mice explored an unaccustomed open field (38×38 cm) for 15 min. Locomotion was analyzed using an ANY-Maze video tracking system (Stoelting Co.), as previously described [17].

### Graded exercise test – ergospirometry

Mice were accustomed to a single-lane treadmill (Panlab LE8710, Harvard Apparatus) at 9 m/min (10 min, slope 5°, and 0.2 mA) with a 24 h interval between each habituation session. The incremental running protocol started at 9 m/min, with an increment of 3 m/min every 2 min at 5° inclination [8,16]. The exercise lasted until running exhaustion, defined by the animal’s inability to leave the electrical grid for five seconds [8,18].

Oxygen uptake 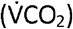 and carbon dioxide production 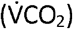 were estimated in a metabolic chamber (Gas Analyzer ML206, 23×5×5 cm, AD Instruments, Harvard) coupled to the treadmill, as previously described [8]. The animals remained in the chamber for 15 min before exercise testing. Atmospheric air (≈21% O_2_, ≈0.03% CO_2_) was renewed at a rate of 120 mL/min, using the same sampling rate for the LASER oxygen sensor (Oxigraf X2004, resolution 0.01%) and infrared carbon dioxide sensor (Servomex Model 15050, resolution 0.1%).

We estimated the running and critical power output in the treadmill based on a standard conversion of the vertical power, body weight, and running speed [8,19]. Running power is the sum (Σ) of all stages of the exercise test, and critical power is the running power performed above 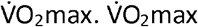 is the maximum capacity to capture (respiratory), transport (cardiovascular), and consume (muscles) oxygen [20]. The respiratory exchange ratio (RER) is the ratio between the amount of carbon dioxide production 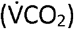 and the consumed oxygen 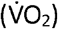 [21].

### Statistics

Data are presented as mean±SD in graphs built using the GraphPad Prism version 5.00 (GraphPad Software, San Diego California USA, www.graphpad.com). Statistical analyzes were performed according to an intention-to-treat principle using StatSoft, Inc. (2007). STATISTICA (data analysis software system), version 13.0. www.statsoft.com. A Student’s t-test was used to evaluate body mass, open field, running power, 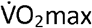, and RER. The evolution of running power and submaximal 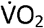 were evaluated by ANOVA for repeated measures followed by a Bonferroni *post hoc* test. The differences were considered significant at P<0.05. Effect sizes (Cohen’s *d*) were calculated for between-group changes in mean differences for open field, running power, and RER, where a Cohen’s *d*=0.2 represents a ‘small’ effect size, 0.5 represents a ‘medium’ effect size, and 0.8 a ‘large’ effect size [22]. Cohen’s η^2^ was used for 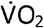 kinetics, defined as small (0.02), medium (0.13) large (0.26) [22].

## RESULTS

The body mass of WT and CD73-KO mice did not differ (t_15_=0.5, Fig.1A), an essential feature since the running power depends on this variable. Locomotion in the open field, either the total distance indicated by the average speed (t_15_=0.75, P=0.4, Fig.1B) or the maximum speed (t_15_=0.16, P=0.87, Fig.1B), was not different between genotypes.

**Fig.1.**
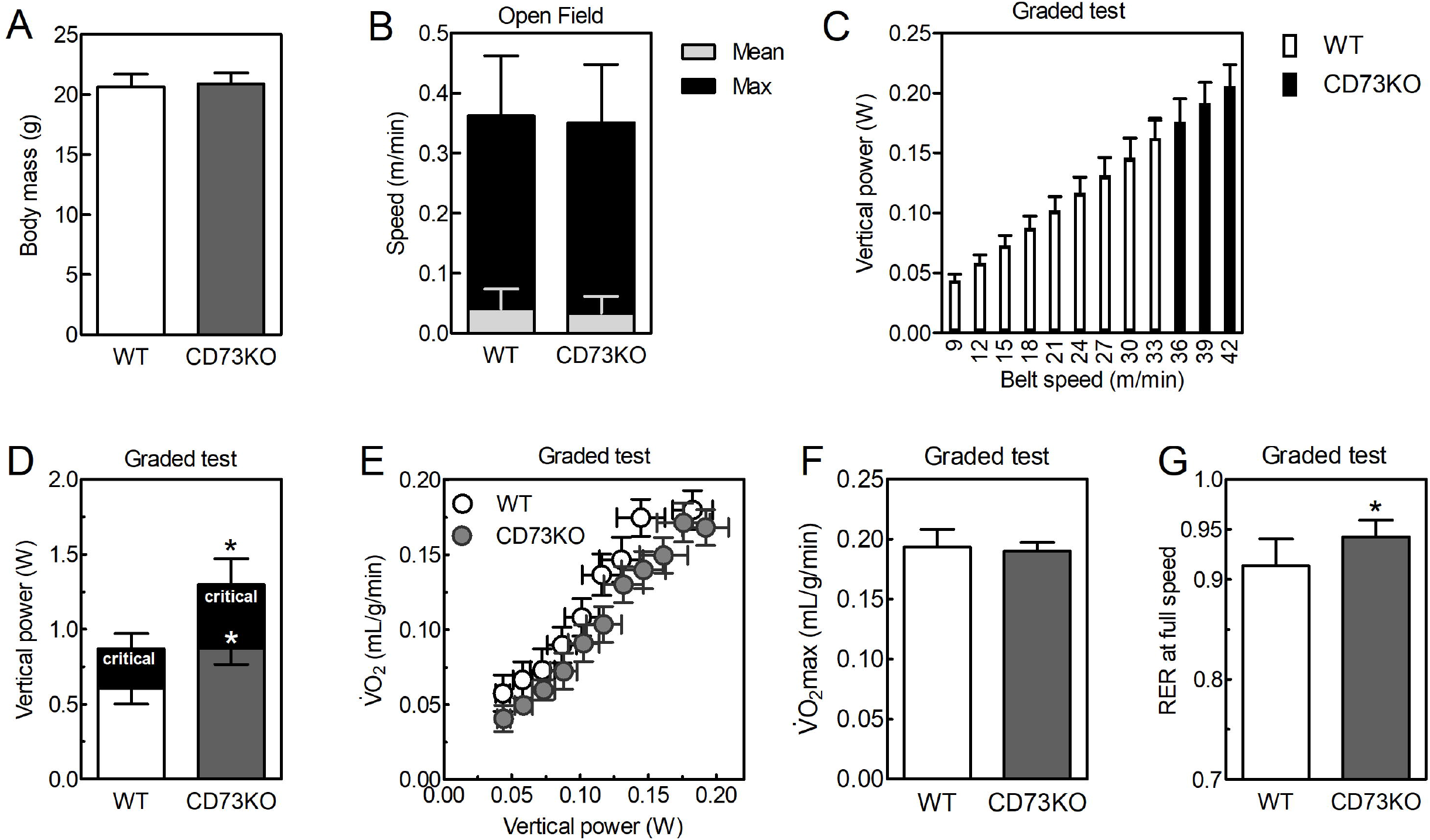
Body mass (A) and locomotion (B) did not differ between wild-type (WT) and CD73 knockout (KO) mice. CD73-KO mice showed greater running power (C and D) and critical power (indicated in red in D), a greater critical power due to the equal 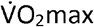 (E and F), and higher carbohydrate consumption indicated by the maximum RER - Respiratory Exchange Ratio (G). Data are described as mean ± SD. N=7-8 animals/group for three independent experiments. *P<0.05 *vs*. WT (Student’s t-test t). 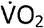 – oxygen consumption.

Running power increased with belt speed acceleration (F_7,56_=30k, P<0.05, η^2^=0.99, Fig.1C), with no differences between genotypes up to 33 m/min (F_7,56_=1.5, P=0.16, Fig.1C); then, only CD73-KO mice continued to run until a maximum belt speed of 42 m/min. Thus, the running power was larger in CD73-KO than WT mice (t_15_=4.2, P<0.05, *d*=0.74, Fig.1D). 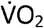 increased (F_7,56_=67, P<0.05, η^2^=0.94, Fig.1E) in line with the intensity of running power, with no difference in submaximal (F_7,56_=0.7, P=0.6, Fig.1E) and maximal 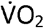 (t_15_=0.25, P=0.8, Fig.1F) between WT and CD73-KO mice.

CD73-KO mice reached a greater critical power (t_15_=5.8, P<0.05, *d*=1.2, Fig.1D) and RER (t_15_=2.4, P<0.05, *d*=0.37, Fig.1G) at the maximum stage of the exercise graded test. Both WT (t_6_=6.9, P<0.05, Fig.1G) and CD73-KO mice (t_6_=9.5, P<0.05, Fig.1G) did not reach the maximum RER value of 1.0.

## DISCUSSION

This study shows that the genetic deletion of CD73 results in an ergogenic profile in mice. Although CD73-KO mice displayed submaximal values of running power and 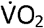 and 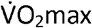 values similar to their WT littermate, CD73-KO mice reached high exercise stages with improved anaerobic power as demonstrated by the greater critical power (large effect) and maximum RER (small effect). This anaerobic power is developed during all-out, short-term exercise above 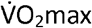 that involves anaerobic muscle metabolism as hinted by the higher maximum RER values characteristic of greater anaerobic consumption of glycolysis.

CD73 controls the formation of adenosine in the periphery and the brain, impacting functions such as motor control, inflammatory responses, adaptive blood pressure, or fatigue. Most of these responses are well-established to be modulated by A_2A_R, in line with the predominant effect of A_2A_R in the ergogenic effect of the non-selective adenosine receptor antagonist, caffeine [8]. A_2A_R is present in blood vessels and their activation triggers reactive vasodilation [23]. Adenosine derived from muscle contraction is responsible for 20-40% of exercise-induced vasodilation [5]. However, adenosine treatment during exercise does not change myocardial and muscle 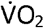 [24,25] and adenosine receptor blockade with 8-phenyltheophylline does not modify tachycardia, mean aortic pressure, coronary blood flow, and myocardial 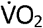 of running dogs [25]. Overall, this suggests a limited impact of vascular A_2A_R on exercise performance, although this remains to be directly tested.

We did not directly assess the alteration of the extracellular levels of adenosine in the blood and different tissues of CD73-KO mice, which is a significant limitation of the present study and a challenge for future studies. However, the combination of current and previous observations also prompts the suggestion that the altered levels of adenosine in the blood of CD73-KO mice [26] might not be the prime contributor to the ergogenic profile CD73-KO mice. CD73-KO mice display few peripheral changes, typified by the lack of altered blood pressure [26], cardiac output and ejection fraction [27], 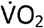 on running wheels during the light and dark phase of circadian rhythm [27], oxygen saturation erythrocytic and pH [27], blood glucose and 2,3-biphosphoglycerate levels [27]. This is in accordance with the presently observed lack of differences in the submaximal and maximum 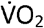 in a graded exercise test between WT and CD73-KO mice. The ergogenic effect resulting from knocking out CD73 cannot also be attributed to motor differences in CD73-KO mice since they do not display alterations of spontaneous locomotion [13,17,28]. Kulesskay et al. [29] observed a hyperlocomotion in CD73-KO mice but attributed this difference to their isolation in cages with running wheels, contrasting with the collective housing in the present study.

The most parsimonious explanation is the possibility that the ergogenic effect of knocking out CD73 might result from a central effect of dampening exercise-induced fatigue. Exercise transiently increases adenosine that can cause sleep and tiredness [4,10,30], whereas caffeine causes arousal and is ergogenic [6,8]. Moreover, the ergogenicity observed in CD73-KO mice phenocopies the previously observed ergogenic profile of forebrain A_2A_R-KO mice [8]. Indeed, CD73 and A_2A_R are functionally coupled in different brain regions [11–13,17], namely, in basal ganglia [12,17], and the hyperlocomotion caused by A_2A_R antagonists is dampened in CD73-KO mice [17].

In conclusion, we now show the critical participation of CD73 in defining the maximum intensity of exercise performance. Although peripheral effects of adenosine cannot be excluded, this effect of knocking out CD73 mimics the impact of knocking out forebrain A_2A_R in controlling fatigue [8], in line with the prominent contribution of central effects of adenosine to control fatigue [6,7].

## Funding

Prémio Maratona da Saúde, CAPES-FCT (039/2014), CNPq (302234/2016-0), LaCaixa Foundation (LCF/PR/HP17/52190001), FCT (POCI-01-0145-FEDER-03127 and UIDB/04539/2020), and ERDF through Centro 2020 (project CENTRO-01-0145-FEDER-000008:BrainHealth 2020 and CENTRO-01-0246-FEDER-000010). A.S.A.Jr is a CNPq fellow (310635/2020-9).

## Conflict of Interest

RAC is a scientific consultant for the Institute for Scientific Information on Coffee. All other authors declare no conflict of interests.

## Ethical approval

Animal experiments were approved by the Ethical Committee of the Center for Neuroscience and Cell Biology (University of Coimbra, ORBEA 138-2016/1507201) and followed the European Union guidelines (2010/63).

## Informed consent

not applicable

## Data availability

The data that support the findings of this study are available from the corresponding author upon reasonable request.

## REFERENCES

1. Borea PA, Varani K, Gessi S, Merighi S, Vincenzi F (2018) The Adenosine Receptors. Springer-Verlag. https://doi.org/10.1007/978-3-319-90808-3

2. Moritz CE, Teixeira BC, Rockenbach L, Reischak-Oliveira A, Casali EA, Battastini AM. (2017) Altered extracellular ATP, ADP, and AMP hydrolysis in blood serum of sedentary individuals after an acute, aerobic, moderate exercise session. Mol Cell Biochem 426: 55–63. https://doi.org/10.1007/s11010-016-2880-1

3. Boussuges A, Rives S, Marlinge M, Chaumet G, Vallée N, Guieu R, Gavarry O (2020) Hyperoxia during exercise: impact on adenosine plasma levels and hemodynamic data. Front Physiol 11: 97. https://doi.org/10.3389/fphys.2020.00097

4. Dworak M, Diel P, Voss S, Hollmann W, Strüder HK (2007) Intense exercise increases adenosine concentrations in rat brain: Implications for a homeostatic sleep drive. Neuroscience 150: 789–795. https://doi.org/10.1016/j.neuroscience.2007.09.062

5. Marshall JM (2007) The roles of adenosine and related substances in exercise hyperaemia. J Physiol 583: 835–845. https://doi.org/10.1113/jphysiol.2007.136416

6. Davis JM, Zhao Z, Stock HS, Mehl KA, Buggy J, Hand GA (2003) Central nervous system effects of caffeine and adenosine on fatigue. Am J Physiol Regul Integr Comp Physiol 284: 399–404. https://doi.org/10.1152/ajpregu.00386.2002

7. Urry E, Landolt HP (2015) Adenosine, caffeine, and performance: from cognitive neuroscience of sleep to sleep pharmacogenetics. Curr Top Behav Neurosci 25: 331–366. https://doi.org/10.1007/7854_2014_274.

8. Aguiar Jr AS, Speck AE, Canas PM, Cunha RA (2020) Neuronal adenosine A_2A_ receptors signal ergogenic effects of caffeine. Sci Rep 10: 13414. https://doi.org/10.1038/s41598-020-69660-1

9. Grgic J, Grgic I, Pickering C, Schoenfeld BJ, Bishop DJ, Virgile A, Pedisic Z (2020) Wake up and smell the coffee: caffeine supplementation and exercise performance – an umbrella review of 21 published meta-analyses. Br J Sports Med 54: 681–688. https://doi.org/10.1136/bjsports-2018-100278

10. Zheng X, Hasegawa H (2016) Administration of caffeine inhibited adenosine receptor agonist-induced decreases in motor performance, thermoregulation, and brain neurotransmitter release in exercising rats. Pharmacol Biochem Behav 140: 82–89. https://doi.org/10.1016/j.pbb.2015.10.019

11. Rebola N, Lujan R, Cunha RA, Mulle C (2008) Adenosine A_2A_ receptors are essential for long-term potentiation of NMDA-EPSCs at hippocampal mossy fiber synapses. Neuron 57: 121–134. https://doi.org/10.1016/j.neuron.2007.11.023

12. Carmo M, Gonçalves FQ, Canas PM, Oses JP, Fernandes FD, Duarte FV, Palmeira CM, Tomé AR, Agostinho P, Andrade GM, Cunha RA (2019) Enhanced ATP release and CD73-mediated adenosine formation sustain adenosine A_2A_ receptor overactivation in a rat model of Parkinson’s disease. Br J Pharmacol 176: 3666–3680. https://doi.org/10.1111/bph.14771

13. Gonçalves FQ, Lopes JP, Silva HB, Lemos C, Silva AC, Gonçalves N, Tomé ÂR, Ferreira SG, Canas PM, Rial D, Agostinho P, Cunha RA (2019) Synaptic and memory dysfunction in a β-amyloid model of early Alzheimer’s disease depends on increased formation of ATP-derived extracellular adenosine. Neurobiol Dis 132: 104570. https://doi.org/10.1016/j.nbd.2019.104570

14. Deaglio S, Dwyer KM, Gao W, Friedman D, Usheva A, Erat A, Chen JF, Enjyoji K, Linden J, Oukka M, Kuchroo VK, Strom TB, Robson SC (2007) Adenosine generation catalyzed by CD39 and CD73 expressed on regulatory T cells mediates immune suppression. J Exp Med 204: 1257–1265. https://doi.org/10.1084/jem.20062512

15. Eltzschig HK, Thompson LF, Karhausen J, Cotta RJ, Ibla JC, Robson SC, Colgan SP (2004) Endogenous adenosine produced during hypoxia attenuates neutrophil accumulation: coordination by extracellular nucleotide metabolism. Blood 104: 3986–3992 https://doi.org/10.1182/blood-2004-06-2066

16. Aguiar AS, Speck AE, Amaral IM, Canas PM, Cunha RA (2018) The exercise sex gap and the impact of the estrous cycle on exercise performance in mice. Sci Rep 8: 10742. https://doi.org/10.1038/s41598-018-29050-0

17. Augusto E, Matos M, Sévigny J, El-Tayeb A, Bynoe MS, Müller CE, Cunha RA, Chen JF (2013) Ecto-5⍰-nucleotidase (CD73)-mediated formation of adenosine is critical for the striatal adenosine A_2A_ receptor functions. J Neurosci 33: 11390–11399. https://doi.org/10.1523/JNEUROSCI.5817-12.2013

18. Ayachi M, Niel R, Momken I, Billat VL, Mille-Hamard L (2016) Validation of a ramp running protocol for determination of the true VO2max in mice. Front Physiol 7: 372. https://doi.org/10.3389/fphys.2016.00372

19. Barbato JC, Koch LG, Darvish A, Cicila GT, Metting PJ, Britton SL (1998) Spectrum of aerobic endurance running performance in eleven inbred strains of rats. J Appl Physiol 85: 530–536. https://doi.org/10.1152/jappl.1998.85.2.530

20. Hill A V, Long CNH, Lupton H (1924) Muscular exercise, lactic acid, and the supply and utilisation of oxygen - Parts ⍰-⍰⍰⍰. Proc R Soc London Ser B 96: 438–475. https://doi.org/10.1098/rspb.1924.0037

21. Naimark A, Wasserman K, McIlroy MB (1964) Continuous measurement of ventilatory exchange ratio during exercise. J Appl Physiol 19: 644–652. https://doi.org/10.1152/jappl.1964.19.4.644

22. Bakeman R (2005) Recommended effect size statistics for repeated measures designs. Behav Res Methods 37: 379–384. https://doi.org/10.3758/BF03192707

23. Johnston-Cox HA, Koupenova M, Ravid K (2012) A_2_ adenosine receptors and vascular pathologies. Arterioscler Thromb Vasc Biol 32: 870–878. https://doi.org/10.1161/ATVBAHA.112.246181

24. Rådegran G, Calbet JAL (2001) Role of adenosine in exercise-induced human skeletal muscle vasodilatation. Acta Physiol Scand 171: 177–185. https://doi.org/10.1046/j.1365-201x.2001.00796.x

25. Bache RJ, Dai XZ, Schwartz JS, Homans DC (1988) Role of adenosine in coronary vasodilation during exercise. Circ Res 62: 846–853. doi: 10.1161/01.res.62.4.846

26. Koszalka P, Özüyaman B, Huo Y, Zernecke A, Flögel U, Braun N, Buchheiser A, Decking UK, Smith ML, Sévigny J, Gear A, Weber AA, Molojavyi A, Ding Z, Weber C, Ley K, Zimmermann H, Gödecke A, Schrader J (2004) Targeted disruption of cd73/ecto-5⍰ nucleotidase alters thromboregulation and augments vascular inflammatory response. Circ Res 95: 814–821. https://doi.org/10.1161/01.RES.0000144796.82787.6f

27. O’Brien WG, Berka V, Tsai AL, Zhao Z, Lee CC (2015) CD73 and AMPD3 deficiency enhance metabolic performance via erythrocyte ATP that decreases hemoglobin oxygen affinity. Sci Rep 5: 13147. https://doi.org/10.1038/srep13147

28. Zlomuzica A, Burghoff S, Schrader J, Dere E (2013) Superior working memory and behavioural habituation but diminished psychomotor coordination in mice lacking the ecto-5⍰-nucleotidase (CD73) gene. Purinergic Signal 9: 175–182. https://doi.org/10.1007/s11302-012-9344-1

29. Kulesskaya N, Võikar V, Peltola M, Yegutkin GG, Salmi M, Jalkanen S, Rauvala H (2013) CD73 is a major regulator of adenosinergic signalling in mouse brain. PLoS One 8: e66896. https://doi.org/10.1371/journal.pone.0066896

30. Nunes EJ, Randall PA, Santerre JL, Given AB, Sager TN, Correa M, Salamone JD (2010) Differential effects of selective adenosine antagonists on the effort-related impairments induced by dopamine D1 and D2 antagonism. Neuroscience 170: 268–280. https://doi.org/10.1016/j.neuroscience.2010.05.068

